# A naïve genome reveals the evolutionary dependence of CTCF functions

**DOI:** 10.64898/2026.11.23.768571

**Authors:** Marta Portela, Alberto Pineda-Santaella, Sergio Camacho-Cabañas, Inmaculada Vázquez-Gutiérrez, María José López-Fernández, Alberto Jiménez-Martín, José M. Santos-Pereira, Alfonso Fernández-Álvarez, Ignacio Maeso

## Abstract

The functions of genome regulatory proteins often emerge through long-term coevolution with their genomic and nuclear environments, complicating the distinction between intrinsic protein properties and context-dependent activities. The DNA-binding protein CTCF is a multifunctional regulator that plays central roles in genome organization and gene regulation across animals. Here, we investigated the behaviour of human CTCF in a deeply heterologous system by expressing it in the budding yeast *Saccharomyces cerevisiae*, a lineage that diverged before the evolution of CTCF itself. Human CTCF was efficiently expressed, localized to the nucleus and associated with yeast chromatin, yet failed to establish robust sequence-specific binding or substantial transcriptional responses. Genome-wide analyses revealed no significant changes in chromatin accessibility, although modest footprinting and nucleosome-positioning effects were detected around predicted CTCF motifs. Together, our results suggest that chromatin association can occur independently of prior evolutionary history, whereas key aspects of CTCF target recognition and regulatory activity depend on long-term genome-protein coevolution.

## INTRODUCTION

The CCCTC-binding factor (CTCF) is a highly conserved zinc-finger protein that plays multiple and essential roles in genome regulation (Corin et al., 2025; Lobanenkov et al., 1990). Since its emergence in the last common ancestor of bilaterian animals (Heger et al., 2012), CTCF performs a diverse array of functions in different animal lineages and has been suggested to play a role in the appearance of complex long-range gene regulation in animals (Acemel et al., 2017; Acemel & Lupiáñez, 2023; Heger et al., 2012; Irimia & Maeso, 2019). In vertebrates, CTCF plays a central role in three-dimensional genome organization (Andreu et al., 2022; Chen et al., 2019; Franke et al., 2021; Niu et al., 2021). Through its interaction with cohesin, CTCF acts as a major architectural determinant of loop extrusion-mediated genome folding, facilitating distal regulatory interactions (Fudenberg et al., 2016; Nora et al., 2017; Rao et al., 2014; Wendt et al., 2008). In addition to these architectural functions, CTCF also participates in transcriptional regulation, insulation, nucleosome positioning, and chromatin state maintenance, highlighting its remarkable functional versatility across cellular contexts (reviewed in Corin et al., 2025; Phillips & Corces, 2009).

Despite its prominent role in 3D chromatin organization in vertebrates, the architectural roles of CTCF in other lineages are much more contentious (Davies et al., 2026; Kaushal et al., 2021; Magri et al., 2026). In particular, recent works have shown that CTCF’s ability to act as an efficient cohesin barrier and to support loop extrusion-based chromatin folding, emerged relatively late during evolution, at the origin of the vertebrate lineage (Astica et al., 2026; Nurislamov et al., 2026). Functional replacement experiments show that vertebrate and non-vertebrate orthologues retain substantial regulatory functions when transferred to a heterologous species, but differ markedly in their ability to support particular architectural functions (Astica et al., 2026; Nurislamov et al., 2026). These findings suggest that the transcriptional and architectural roles of CTCF are evolutionarily separable, with some of these functions being most likely ancestral and intrinsic to the protein, whereas others depend on evolutionary adaptations that occurred within specific genomic and cellular contexts.

However, previous cross-species approaches have largely been restricted to organisms that possess endogenous CTCF orthologues. Even when CTCF proteins are experimentally transferred between species, they are introduced into genomes that share a long evolutionary history with CTCF and in which DNA sequence composition, chromatin organization, and regulatory networks have been shaped by its presence. Thus, it remains difficult to determine which features of CTCF emerge directly from its biochemical properties and which require extensive coevolution between the protein and its genomic substrate.

The budding yeast *Saccharomyces cerevisiae* offers an opportunity to address this. Fungi diverged from the animal lineage before the emergence of CTCF. Therefore, the yeast genome evolved in the complete absence of CTCF-mediated regulation: its sequence composition, chromatin organization and gene regulatory networks have not undergone selection in response to this factor. Expression of CTCF in yeast therefore offers an opportunity to evaluate which aspects of CTCF behaviour can emerge in a naïve eukaryotic system that has never coevolved with the protein.

Through the expression of human CTCF in the deeply heterologous context of *S. cerevisiae,* here we combine transcriptomic, chromatin-binding, and chromatin accessibility analyses to investigate which aspects of CTCF behaviours are independent from its long evolutionary history within animal genomes.

## RESULTS

### Human CTCF is expressed, localized to the nucleus and bound to chromatin in *S. cerevisiae*

To express human CTCF (hCTCF) in a genome that has never been exposed to it, we cloned the full-length hCTCF coding sequence into the centromeric vector pRS416 under the control of the galactose-inducible *GAL1* promoter, and transformed *S. cerevisiae* BY4741 strain with this construct (+CTCF) or with the corresponding empty vector (control pØ) (Fig. 1A). Expression of a vertebrate architectural protein might be expected to interfere with chromosome biology, yet ten-fold serial dilution assays of three independent transformants per construct revealed no growth defect: +CTCF and control pØ cells grew indistinguishably on rich medium (YPDA) and on selective medium (YNB–U) under both repressing (raffinose) and inducing (raffinose plus galactose) conditions, and across the whole range of temperatures tested, from 25 to 36°C (Fig. 1B; Supplementary fig. 1). hCTCF is therefore tolerated in budding yeast without a detectable fitness cost.

**Figure 1.**
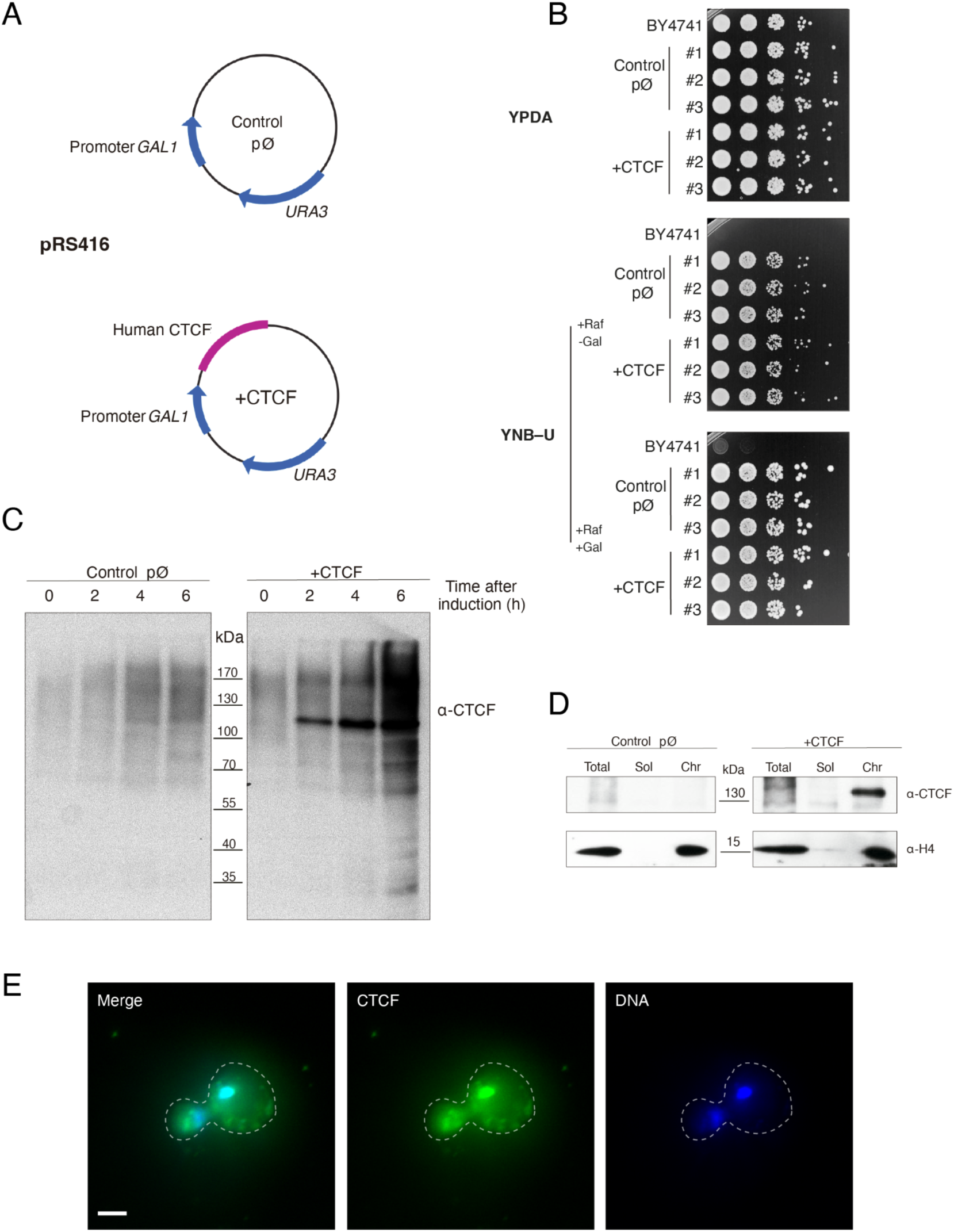
Human CTCF is expressed, imported into the nucleus and bound to chromatin in *S. cerevisiae*. **(A)** Constructs used in this study: the centromeric plasmid pRS416 carrying *URA3* and the *GAL1* promoter, either empty (control pØ) or driving the full-length human CTCF coding sequence (+CTCF). **(B)** Ten-fold serial dilution growth assays of the parental strain BY4741 and of three independent transformants (#1–#3) per construct, spotted on rich medium (YPDA) and on selective medium lacking uracil (YNB–U) under repressing (+Raf −Gal) and inducing (+Raf +Gal) conditions, and incubated at 28 °C for two days (YPDA) or four days (YNB–U). As expected for a *ura3Δ0* strain, BY4741 does not grow on YNB–U. The complete temperature series is shown in supplementary figure S1. **(C)** Western blot of total protein extracts obtained 0, 2, 4 and 6 h after the addition of 2% galactose, probed with anti-CTCF. The hCTCF-specific band migrates at the expected ∼130 kDa; molecular weight markers (kDa) are indicated. **(D)** Western blot of total (Total), soluble (Sol) and chromatin-enriched (Chr) fractions prepared 4 h after induction, probed with anti-CTCF and anti-histone H4; H4 marks the chromatin fraction. Both panels derive from the same membrane, probed sequentially with anti-CTCF and anti-histone H4. **(E)** Immunofluorescence of an induced +CTCF cell showing the merge of both channels, hCTCF (green) and DNA (blue). Scale bar, 2 µm.

We next verified that the protein accumulated as a full-length product. Western blot of total extracts collected 0, 2, 4 and 6 h after the addition of 2% galactose detected an hCTCF-specific band of the expected apparent molecular weight (∼130 kDa) exclusively in +CTCF cells. The band was already evident 2 h after induction and increased thereafter (Fig. 1C). Lower-molecular-weight species became apparent at 6 h, consistent with partial degradation of the protein upon prolonged induction, and all subsequent experiments were therefore performed 4 h after induction.

To determine whether hCTCF binds to yeast chromatin, we fractionated spheroplast lysates into soluble and chromatin-enriched fractions 4 h after induction. hCTCF was recovered almost exclusively in the chromatin fraction, defined by the presence of histone H4, with only a minor fraction remaining soluble (Fig. 1D). Consistent with this, immunofluorescence of induced cells showed hCTCF signal concentrated over the DAPI-stained nuclear mass rather than distributed throughout the cytoplasm (Fig. 1E). Thus, hCTCF produced in a fungal cell is correctly imported into the nucleus and stably associated with chromatin, in the absence of any CTCF-specific partner and with a genome that has never been selected for its binding.

### Limited transcriptional consequences of heterologous hCTCF expression

Given the observed binding of hCTCF to the yeast genome and its potential effects on chromatin organization, we sought to determine whether these interactions were associated with changes in gene expression. To that end we performed an RNA-seq and compared hCTCF-expressing and control yeast cells. Differential gene expression analysis revealed only 18 significantly upregulated genes, most of which were mitochondrial genes (Fig. 2A, Supp. Fig. 2A). To investigate whether hCTCF expression affected specific biological processes despite these limited transcriptional changes, we performed gene set enrichment analysis (GSEA). This analysis revealed that upregulated gene sets were predominantly associated with ion transport, whereas downregulated gene sets were mainly related to ribosomal homeostasis (Fig. 2B; Supp. Table 1).

**Figure 2.**
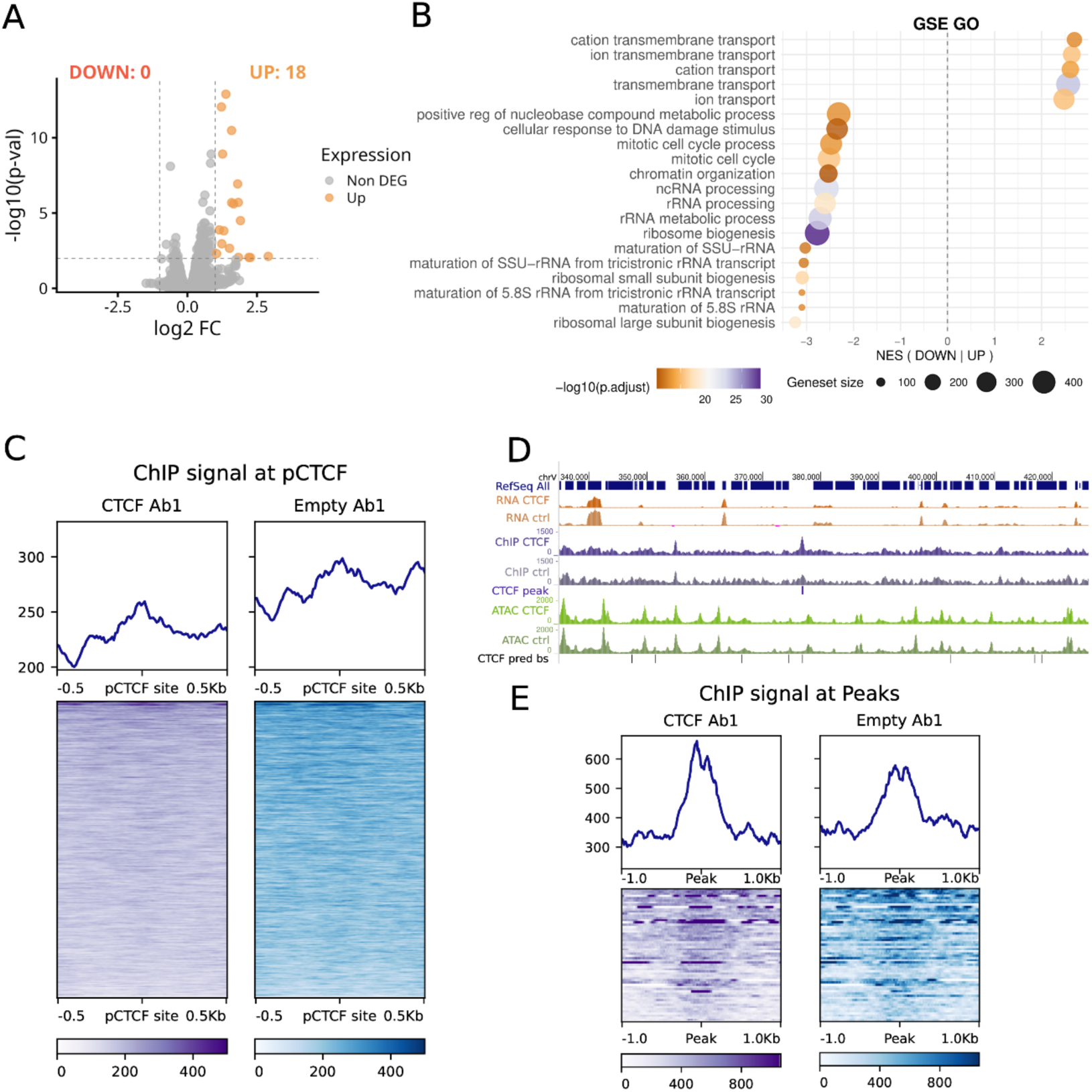
Gene expression changes after expression of hCTCF and ChIPmentation profiles. **(A)** Volcano plots showing differential gene expression between hCTCF-expressing yeast and control RNA-seq datasets. 18 genes have been found to be significantly upregulated at |log2FC| > 1 and pvalue < 0.01. **(B)** Gene set enrichment analysis (GSEA) of Gene Ontology (GO) biological processes (BP), showing the top 20 enriched terms. Dot size represents the number of genes associated with each term, while color indicates the statistical significance of enrichment. **(C)** ChiPmentation profile across a 1 kb region centered on the predicted hCTCF binding sites in hCTCF-expressing and control yeast. **(D)** Screenshot of *S. cerevisiae* genome browser, displaying RNA-seq expression signal (orange), ChIPmentation signal of CTCF (purple), CTCF called peaks (dark purple), ATAC-seq signal (green) and fimo-predicted hCTCF binding sites (black). **(E)** ChIPmentation signal profiles across 2 kb centered at hCTCF ChIPmentation peaks but absent in control yeast, comparing hCTCF-expressing and control conditions.

These modest transcriptional changes, that oscillate at very low foldchanges, are consistent with the survival results obtained. They also correlate with previous studies showing that global CTCF depletion in mammalian cells generally results in limited and context-dependent effects on gene expression, despite substantial alterations in chromatin architecture (Nora et al., 2017; Zuin et al., 2014). Moreover, CTCF-mediated transcriptional regulation has been shown to depend on specific genomic contexts, including promoter-proximal binding that facilitates enhancer-dependent gene activation (Kubo et al., 2021).

### hCTCF binding in yeast genome lacks specificity

We therefore sought to determine whether the transcriptional changes observed upon hCTCF expression in *S. cerevisiae* could be associated with its direct binding to the yeast genome or instead reflected indirect effects of heterologous protein expression. To address this question, we performed a ChIPmentation assay (Schmidl et al., 2015), to characterize the genome-wide binding profile of hCTCF. We first examined whether CTCF preferentially occupied genomic regions containing predicted hCTCF-binding motifs in the yeast genome. We identified 1,702 motif occurrences with a p-value less than 0.0001 and the ChIPmentation signal at these sites showed no detectable enrichment in hCTCF-expressing yeast compared with yeast carrying the control vector (Fig. 2C). We conducted these experiments in parallel with two different antibodies against hCTCF to avoid any biochemical bias. Similarly, we observed no preferential accumulation of hCTCF signal around transcription start sites (TSSs), in contrast to Pol II distribution (Supp. Fig. 2D), suggesting that hCTCF does not exhibit a general preference for promoter-proximal regions in the yeast genome.

Nevertheless, visual inspection of the ChIPmentation profiles revealed several discrete regions displaying increased signal in hCTCF-expressing samples (Fig. 2D). To systematically characterize these potential binding events, we performed peak calling and identified 54 peaks exclusively detected in hCTCF-expressing samples. However, these peaks showed limited overlap between the antibodies used for hCTCF detection (Supp. Fig. 2C) and exhibited only modest signal enrichment relative to control samples (Fig. 2E). Furthermore, *de novo* motif enrichment analysis failed to identify any significantly positionally enriched sequence motifs or motifs resembling the canonical hCTCF-binding motif (Supp. Fig. 2E).

### Chromatin accessibility and nucleosome organization upon ectopic hCTCF expression

Even if hCTCF is able to bind the yeast chromatin, as we saw above (Fig. 1D), the absence of a robust sequence-specific hCTCF binding raises the possibility that nucleosome occupancy or local chromatin accessibility may limit its interaction with the yeast genome. Conversely, CTCF itself has been implicated in shaping local nucleosome organization, suggesting that its expression could influence chromatin structure (Fu et al., 2008; Owens et al., 2019). To explore these possibilities, we performed ATAC-seq to investigate changes in chromatin accessibility and nucleosome positioning following hCTCF expression in yeast.

ATAC-seq analysis revealed that treated yeast exhibited a modest increase in overall chromatin accessibility, as reflected by slightly higher ATAC-seq signal and a greater number of called peaks, 4,149 versus 3,868 in control yeast cells (Fig 3A, Supp. Fig. 3A and 3B). However, differential accessibility analysis did not identify any significant differentially accessible regions (Supp. Fig. 3C). Furthermore, no differences in accessibility were observed at the candidate CTCF-associated regions identified by ChIPmentation (Supp. Fig. 3D), nor was an increased CTCF ChIPmentation signal detected at ATAC-seq peaks (Supplementary Fig. 3E).

**Figure 3.**
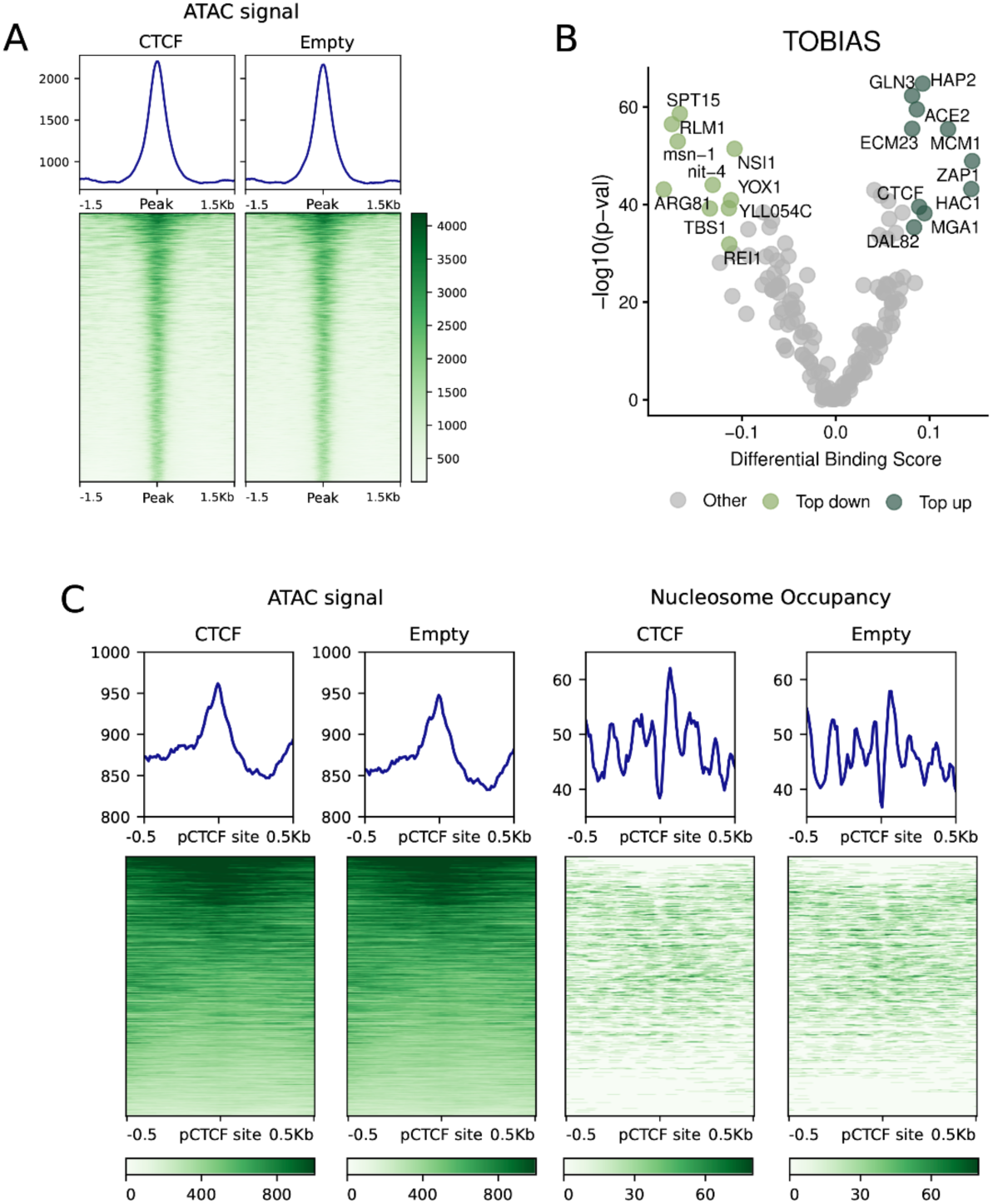
Chromatin accessibility after the expression of hCTCF. **(A)** ATAC-seq signal profiles across a 3 kb region centered on the global ATAC-seq peakset between the hCTCF-expressing yeast and the control. **(B)** Volcano plot displaying the differential motif occupancy scores reflecting global changes in TF binding in the global ATAC-seq peakset between the hCTCF-expressing yeast and the control from TOBIAS footprinting analysis. Top 10 motifs, up and down, are coloured. **(C)** ATAC-seq signal profiles (left) and nucleosome occupancy (right) across a 1 kb region centered on the predicted hCTCF binding sites in hCTCF-expressing and control yeast.

To further investigate whether these subtle accessibility differences were accompanied by changes in transcription factor occupancy, we performed ATAC-seq footprinting analysis using TOBIAS (Bentsen et al., 2020). Interestingly, the CTCF motif was among the top 10 showing increased footprinting signals in hCTCF-expressing yeast compared with controls, although these differences were modest (Fig. 3B; Supp. Table 3). Finally, since CTCF binding has been associated with the organization of flanking nucleosomes in mammalian cells (Fu et al., 2008; Owens et al., 2019), we next examined nucleosome positioning around predicted CTCF-binding sites in the yeast genome. This analysis revealed a modest increase in the nucleosome-associated signal surrounding these sites in hCTCF-expressing yeast (Fig. 3C), suggesting subtle changes in local chromatin organization despite the absence of robust CTCF binding detected by ChIPmentation.

## DISCUSSION

In this work we have deepened our understanding of the ability of CTCF to interact in a heterologous context, the genome of *S. cerevisiae*, a lineage that diverged before the emergence of CTCF itself and whose genome therefore evolved in complete absence of CTCF-mediated regulation. Our results reveal a striking contrast between chromatin association and functional regulatory activity. hCTCF is efficiently expressed in yeast, accumulates in the nucleus and successfully interacts with fungal chromatin, despite the absence of known metazoan-specific partners or a shared evolutionary history between CTCF and its new ectopic genomic substrate.

However, despite the successful binding of hCTCF to the yeast genome, we did not observe significant changes in chromatin accessibility or robust sequence-specific binding. There was no preferential occupancy of predicted CTCF motifs or promoter-proximal regions, and limited transcriptional changes. Our findings contrast with recent works performing CTCF functional replacements across different bilaterian species (Astica et al., 2026; Nurislamov et al., 2026). These experiments revealed the presence of lineage-specific architectural functions, but in all cases, the different CTCF orthologs exhibited robust sequence-specific binding, clearly showing the distinctive motif preferences of each lineage. Together, these results suggest that in a naïve eukaryotic context, CTCF chromatin association alone may be insufficient to induce chromatin opening and establish the robust canonical binding and regulatory behaviours characteristic of animal genomes. Although CTCF has been proposed to exhibit pioneer-like properties in certain cellular contexts, its ability to function as a pioneer transcription factor remains debated (Tajmul et al., 2025). Notably, a Gal4–CTCF fusion was reported to act as an insulator in budding yeast when artificially tethered to a telomeric reporter (Defossez & Gilson, 2002). The contrast with our genome-wide data suggests that CTCF can locally interfere with chromatin spreading once recruited to a defined position, but does not autonomously locate and act on its motifs in a genome that has not coevolved with it. These findings raise the possibility that CTCF does not function as an autonomous pioneer factor in this heterologous system, and that its chromatin-remodelling activity may depend on additional cofactors or specific chromatin contexts that are absent in yeast.

Similar constraints have recently been reported for multiple human TFs expressed in budding yeast, where 30% of the tested TFs displayed weak or no binding signal (Bugis et al., 2026). Motif length may contribute to this: the 178 TFs encoded in budding yeast have binding motifs of 7.92 bp on average, whereas the three motifs reported for hCTCF average 26.33 bp (15– 33 bp) (Nakahashi et al., 2013), and the human TFs that failed to bind yeast chromatin in Bugis et al. recognise motifs of 11–14 bp. The compact nature of the yeast genome, with far fewer flexible, loosely occupied intergenic regions than animal genomes, may act as an additional constraint for TFs with different pioneer capacities.

At the same time, our data do not support a complete absence of chromatin-related activity. Although no significant differences in chromatin accessibility were detected, ATAC-seq analyses ranked the CTCF motif among the highest differential footprinting scores –although these differences were small— together with subtle alterations in local nucleosome organization surrounding these regions. While these effects should be interpreted cautiously, they are consistent with previous observations linking CTCF to local nucleosome positioning (Nurislamov et al., 2026; Owens et al., 2019; Tajmul et al., 2025) and raise the possibility that some chromatin-interacting properties of the protein can emerge even in a deeply heterologous context.

In sum, our findings illustrate how the functional properties of genome regulators cannot always be understood as intrinsic attributes of individual proteins, but often emerge from long-term evolutionary interactions between regulatory factors and their genomes.

## METHODS

### Yeast strains, plasmids and growth conditions

All experiments were performed in the *S. cerevisiae* BY4741 background (*MATa his3Δ1 leu2Δ0 met15Δ0 ura3Δ0*). The full-length human CTCF coding sequence was cloned into the centromeric plasmid pRS416 under the control of the galactose-inducible GAL1 promoter; the corresponding empty vector was used as a control. Both plasmids were introduced into BY4741 using the lithium acetate/single-stranded carrier DNA/polyethylene glycol method (Gietz & Woods, 2002), yielding strains AFA3252 (+CTCF) and AFA3246 (control pØ). Transformants were selected and maintained on synthetic complete medium lacking uracil (SC–U). Unless otherwise stated, cells were grown at 28 °C with shaking in SC–U containing 2% raffinose as the carbon source. Rich medium (YPDA) and synthetic medium lacking uracil (YNB–U) were used where indicated.

### Galactose induction of hCTCF expression

Expression from the GAL1 promoter was induced as follows. A 10 mL preculture was grown in SC–U with 2% raffinose for approximately 12 h and used to inoculate a 50 mL subculture, which was grown overnight under the same conditions and then diluted from OD_600_ ≈ 1 to OD_600_ ≈ 0.3. When cultures reached OD_600_ = 0.5, galactose was added to a final concentration of 2% to induce expression; non-induced controls were maintained in 2% raffinose. For the induction time course, samples equivalent to 10 mL at OD_600_ = 0.5 were collected at 0, 2, 4 and 6 h after the addition of galactose. For all other experiments, cells were harvested 4 h after induction.

### Serial dilution growth assays

Growth was assessed by ten-fold serial dilution spotting. Three independent transformants of each construct, together with the untransformed parental strain BY4741, were grown to exponential phase, adjusted to the same OD_600_ and spotted as ten-fold serial dilutions onto rich medium (YPDA) and onto synthetic medium lacking uracil (YNB–U) containing 2% raffinose, either without galactose (repressing, +Raf −Gal) or with 2% galactose (inducing, +Raf +Gal). Plates were incubated at 25, 28, 32 and 36 °C and imaged after two days (YPDA) or four days (YNB–U). Two independent biological replicates were performed. The 28 °C plates are shown in Figure 1B and the complete temperature series in Supplementary Fig. 1.

### Chromatin fractionation and Western Blot

*S. cerevisiae* cells containing either the empty vector (Control pØ) or the galactose-induced CTCF vector (+CTCF) were grown in synthetic complete medium lacking uracil (SC-U) with raffinose 2% as the carbon source. 4 hours after the addition of galactose 2%, cells were obtained from 50 ml of exponential growing cultures at OD_600_ = 0.5 at 28°C. Cells were collected by centrifugation at 4°C. After incubation with cell wall-relaxing buffers, they were incubated at 30°C with 50 µl of zymolyase 20 T for 1 hour. Once the spheroplasts were generated, they were lysed and the total, soluble and chromatin fractions were separated by centrifugations. Proteins were detected with an anti-CTCF antibody (Sigma-Aldrich, 07-729) (1:5000) and an anti-histone H4 antibody (Abcam, ab10158) (1:3000). The secondary antibody used was horseradish peroxidase (HRP)-coupled anti-rabbit IgG (Sigma-Aldrich, A0545). The visualisation was performed using the WesternBright Sirius detection kit (Advansta) in a Chemidoc MP Imaging System (Biorad). Images processing for representation was performed using Adobe Photoshop CS5 Extended.

### Immunostaining

Immunofluorescence was performed following the protocol described by Silver (2009), with minor modifications. Briefly, cells were obtained from a 5 mL preculture of the BY4741-CTCF strain grown in YNGal-ura medium. The protocol was modified by introducing an additional 1h incubation in BSA-PBS. CTCF was detected using an anti-CTCF antibody (D31H2 XP, 1:800 dilution), followed by an Alexa Fluor 488-conjugated secondary antibody (1:1000 dilution). Fluorescence images were acquired using a DeltaVision Restoration Microscopy System equipped with a 100× oil-immersion objective.

### RNA extraction and library preparation

Total RNA was extracted from exponentially growing *Saccharomyces cerevisiae* cultures using the RNeasy Mini Kit (QIAGEN). Cells were mechanically disrupted using glass beads in RLT buffer supplemented with β-mercaptoethanol. RNA was purified using silica-based spin columns, including on-column DNase I treatment to remove genomic DNA contamination. RNA concentration and purity were assessed using a NanoDrop spectrophotometer. Ribosomal RNA was depleted using the NEBNext RNA Depletion Core Reagent Set (New England Biolabs, Cat. No. E7865L), and libraries were subsequently prepared using the TruSeq Stranded mRNA Kit (Illumina, Cat. No. 20020594).

### ATAC-seq library preparation

ATAC-seq libraries were prepared following the protocol described by Hendrickson et al., (2018). Briefly, 1–5 million yeast cells were treated with zymolyase to generate spheroplasts, followed by transposition using Tn5 transposase (Nextera DNA Sample Preparation Kit, Illumina). Tagmented DNA was purified, PCR-amplified using indexed primers and size-selected using AMPure XP beads.

### ChIPmentation library preparation

ChIPmentation libraries were prepared following a modified version of the protocol described by Schmidl et al. (2015). Briefly, yeast cultures at an OD_600_≈1 were fixed with 1% formaldehyde for 15 min at room temperature and quenched with 125 mM glycine for 5 min. Cells were washed twice with cold PBS, collected by centrifugation and stored at −80°C until further processing. Chromatin extraction was performed using glass bead mechanical disruption in ChIP lysis buffer, followed by sonication (20% duty cycle, 75 W peak incident power, 200 cycles per burst, 30 min) to generate fragmented chromatin.

For immunoprecipitation, sonicated chromatin was incubated overnight at 4°C with 1–5 μg of specific antibody. Immune complexes were recovered using Protein G Dynabeads (Thermo Fisher Scientific) and subjected to sequential washes with lysis buffer, high-salt lysis buffer (500 mM NaCl) and LiCl detergent wash buffer. Bead-bound chromatin was subsequently tagmented by incubation with Tn5 transposase (Illumina Nextera DNA Sample Preparation Kit) in tagmentation reaction buffer (10 mM Tris-HCl pH 8.0, 5 mM MgCl_2_, 10% dimethylformamide) at 37°C for 1 min. Following additional washes, DNA was eluted, reverse cross-linked in the presence of NaCl and Proteinase K, and purified using MinElute PCR Purification columns (QIAGEN).

### Libraries quality and sequencing

Library concentrations and fragment size distributions were assessed using a Qubit 3.0 fluorometer (Thermo Fisher Scientific) and an Agilent 2100 Bioanalyzer with the High Sensitivity DNA Kit (Agilent Technologies, Cat. No. 5067-4626), respectively. Libraries were sequenced on an Illumina NextSeq 500 platform.

### Genomic data processing

Quality control of raw RNA-seq, ATAC-seq and ChIP-seq data was performed using FastQC v0.12.0 (Andrews, 2010) to evaluate basic quality metrics, followed by trimming with trimGalore v0.6.10 (Krueger et al., 2023) to remove adapter sequences and low-quality bases. The following parameters were used for trimming: −2colour 20 --stringency 3 and clipping 3 bases in the 3’ end and 10 bases in the 5’ end.

Bowtie2 v2.5.0 (Langmead et al., 2019) was used to align the trimmed reads to the reference genome SacCer3 with the following options -local --very-sensitive-local --no-unal --no-mixed - -no-discordant --dovetail --phred33 -I 10 -X 500. PCR duplicates were removed with picard (Xu et al., 2012). Expected plasmid expression was corroborated by aligning the reads to the plasmid sequence.

Nucleosome positioning was inferred from ATAC-seq data using deNOPA (Xu et al., 2022). ATAC-seq bam files were used to identify nucleosome-associated regions, identifying both nucleosome free regions (NFR) and nucleosome positions, with the number of sequencing fragments intersected. These regions were transformed into continuous data through bamCoverage.

### Differential expression analysis

Following alignment, raw read counts for gene and genomic peaks were quantified using featureCounts with the --countReadPairs option. Differential expression analysis was carried out using DESeq2 v1.46.0 (Love et al., 2014), where differentially expressed genes were considered with a pval<0.01 and a minimum log2FC of +/-1, and plots were generated using ggplot2 v3.5.1 (Villanueva & Chen, 2019). Principal component analysis (PCA) of RNA-seq, ChIPmentarion and ATAC-seq samples was performed with normalized counts of the 500 most variable genes with the R function plotPCA. For the plot representation of the ChIPmentation samples, in order to correct the batch effect, we used ComBat-seq (Zhang et al., 2020). Gene Set Enrichment (GSE) analysis was performed over Biological Processes using the ClusterProfiler package v4.2.0 (Wu et al., 2021) and a p-value cut-off of 0.05.

### Peak calling

Peak calling was performed using MACs2 v2.2.9.1 (Zhang et al., 2008) with experiment-specific parameters. For ChIPmentation data, peaks were called using --nomodel --extsize 200 -p 1e-3, whereas ATAC-seq peaks were identified using --nomodel --extsize 50 -q 0.01.

For both experiments we later measured consistency across biological replicates with the use of IDR v2.0.2 (Irreproducible Discovery Rate) peak calling with default parameters (Li et al., 2011). To identify CTCF peaks and accessible chromatin regions specific to each condition and absent from control samples, genomic intervals were compared using the BEDTools multiinter algorithm v2.31.0 (Quinlan & Hall, 2010).

To explore the signal distribution across given regions (peaks, TSS etc) we plotted continuous data with deepTools package v4.0.0 (Ramírez et al., 2014).

### Motif analysis

To identify putative binding sites for CTCF in the *S. cerevisiae* genome we used FIMO from the MEME Suite (Grant et al., 2011). SacCer3 genome was scanned for the most common human CTCF matrix (MA0139) and individual motif matches were identified based on their statistical significance, following default thresholds.

De novo motif discovery was performed on CTCF called peaks from the ChIPmentation experiments using GimmeMotifs v0.18.4 (van Heeringen & Veenstra, 2011)(--denovo -f 0.4). Then, to assess if the obtained motifs were similar to the hCTCF motif, we run PWMSimilarity with Pearson correlation from TFBSTools (Tan & Lenhard, 2016).

TF binding activity was assessed from ATAC-seq data using BINDetect, implemented in the TOBIAS framework v0.17.5 (Bentsen et al., 2020). BINDetect was applied to the ATAC-seq samples using a custom motif collection comprising 181 fungal TF motifs together with 3 CTCF motifs (MA0139, MA1929 and MA1930). Differential TF binding activity between conditions was determined from TOBIAS footprint scores at predicted motif sites.

## DATA AVAILABILITY

All sequencing data is available at the European Nucleotide Archive (ENA) database repository under the study ID PRJEB129517.

## ACKNOWLEDGEMENTS

We thank Pedro A. San-Segundo (IBFG) for sharing the chromatin fractionation protocol and for providing antibodies, Pedro M. Rocha for advice regarding CTCF antibodies and Mar Sánchez (Genomics Facility, IBFG) for library preparation and for her support with all sequencing work. IM was supported by grants PID2021-128728NB-I00 funded by MICIU/AEI/10.13039/501100011033 and by “ERDF/EU” and PID2025-174555NB-I00 funded by MICIU/AEI/10.13039/501100011033. AF-A is funded by PID2021-127232NB-I00 and PID2024-162761NB-I00 funded by the MCIN/ AEI /10.13039/501100011033 and by the “ERDF/EU”. JMS-P was funded by an “Emergia” grant EMC21_00188 from Junta de Andalucía; grant PID2023-147172NA-I00 by MICIU/AEI/10.13039/501100011033 and by “ERDF/EU”; and grant CNS2023-143945 by MICIU/AEI/10.13039/501100011033 and by “European Union NextGenerationEU/PRTR”. AF-A and IM were funded by a “INTERCAMBIOS CIENTÍFICOS LIFEHUB Jump-Start” grant LifeHUB3_01, funded by LifeHUB.CSIC (PIE 202120E047-Conexiones-Life). MJL-P was a recipient of “Ayudas para contratos predoctorales” PRE2019-087394 funded by MICIU/AEI/10.13039/501100011033 and by “ESF+”. IV-G was a recipient of a “JAE-Intro” fellowship JAEINT_19_02145 funded by “Programa JAE” of “Consejo Superior de Investigaciones Científicas” (CSIC).

## SUPPLEMENTARY MATTERIAL

### Supplementary Figures

**Supplementary Figure 1.** Expression of human CTCF does not affect yeast growth across temperatures, and the protein accumulates in the nucleus.

**Supplementary Figure 2.** Gene expression changes after expression of hCTCF and ChIPmentation profiles.

**Supplementary Figure 3.** Chromatin accessibility after the expression of hCTCF.

### Supplementary Tables

**Supplementary Table 1.** RNA-seq and ATAC-seq differential analyses

**Supplementary Table 2.** ATAC-seq and ChIPmentation analyses

---

**Supplementary Figure 1.**
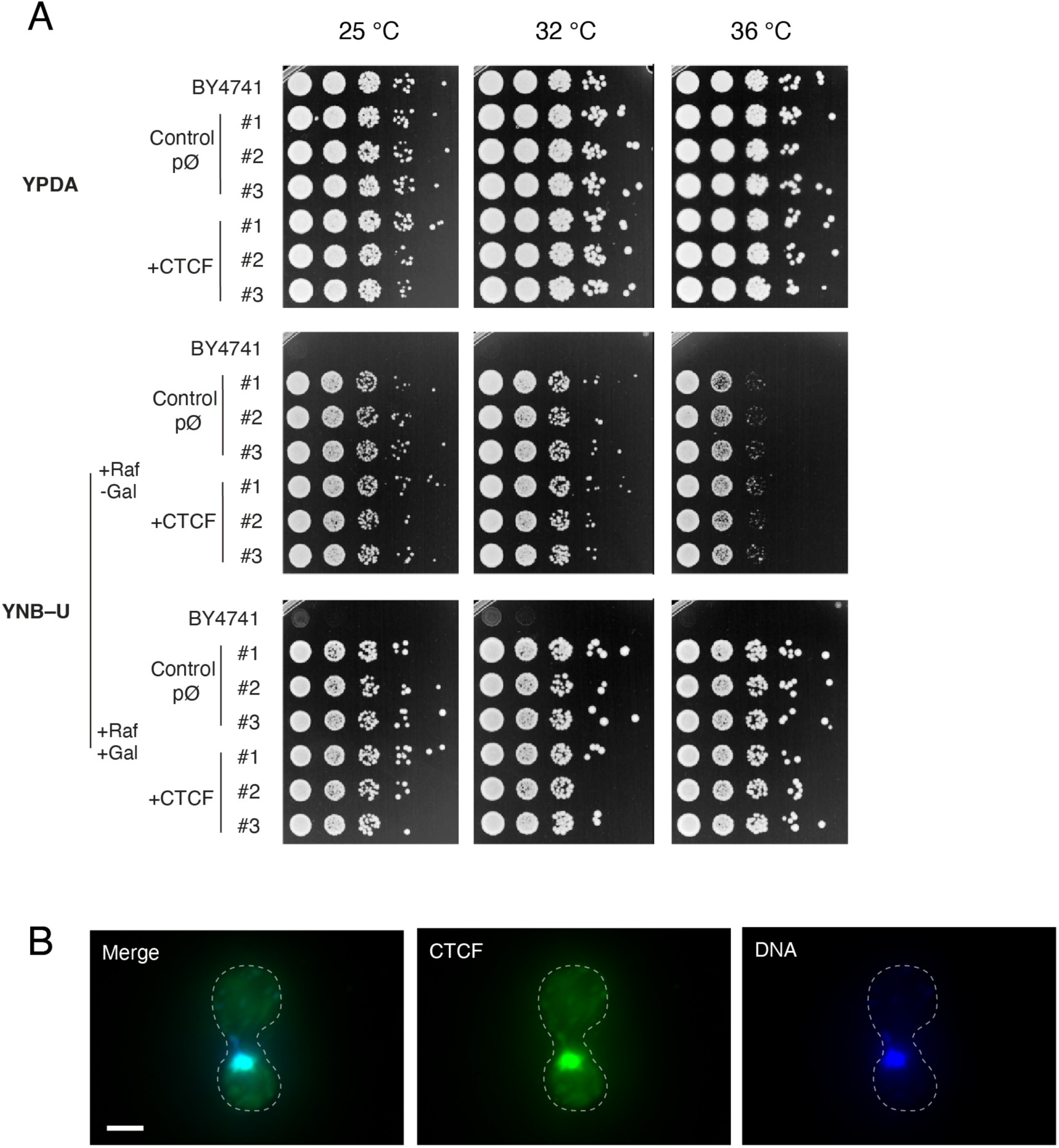
Expression of human CTCF does not affect yeast growth across temperatures, and the protein accumulates in the nucleus. **(A)** Ten-fold serial dilution growth assays of the parental strain BY4741 and of three independent transformants (#1–#3) carrying either the empty vector (control pØ) or the human CTCF construct (+CTCF), spotted on rich medium (YPDA) and on selective medium lacking uracil (YNB–U) under repressing (+Raf −Gal) and inducing (+Raf +Gal) conditions, and incubated at 25, 32 and 36 °C. Plates were imaged after two days (YPDA) or four days (YNB–U). The 28 °C series is shown in Figure 1B. As expected for a *ura3Δ0* strain, BY4741 does not grow on YNB–U. **(B)** Immunofluorescence of a representative +CTCF cell, showing the merge of both channels, hCTCF (green) and DNA (blue). Dashed lines indicate the cell outline. Scale bar, 2 µm.

**Supplementary Figure 2.**
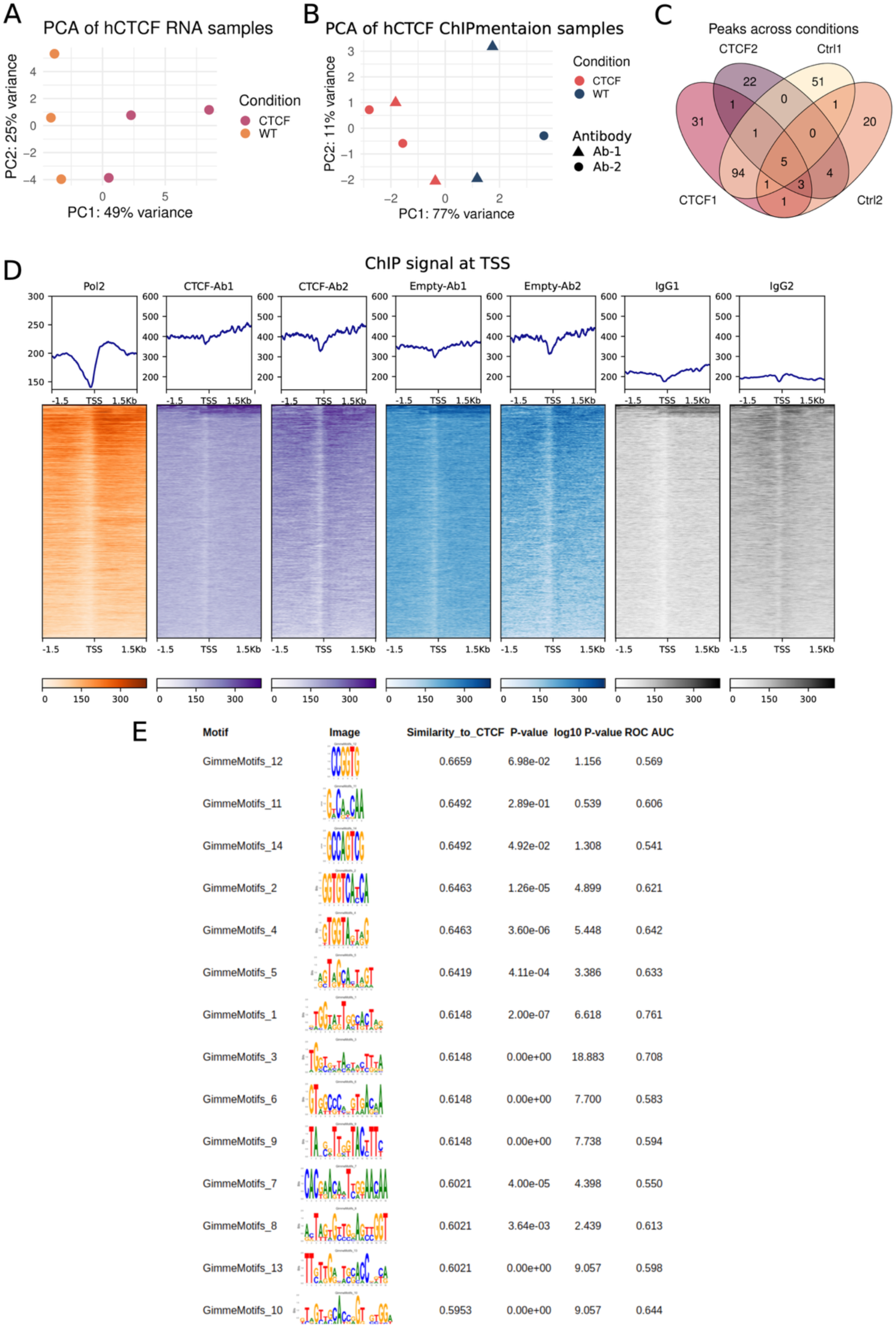
Gene expression changes after expression of hCTCF and ChIPmentation profiles. **(A)** Principal component analysis of RNA-seq transcriptomic profiles from hCTCF expressing yeasts and control yeasts. Each point represents one biological replicate (n=3 per group). PC1 and PC2 explain 49% and 25% of the variance, respectively. **(B)** Principal component analysis of ChIPmentation profiles from hCTCF expressing yeasts and control yeasts. Each point represents one biological replicate (n=4 per group). The antibodies used are defined by the shape of the replicate. PC1 and PC2 explain 77% and 11% of the variance, respectively. **(C)** Venn diagram showing the overlap of the ChIPmentation peaks across experimental conditions. Numbers 1 and 2 denote the antibodies used for ChIPmentation. **(D)** ChIPmentation signal profiles for RNA polymerase II (Pol II), CTCF, and IgG across a 3kb region centered at gene transcription start sites (TSS). **(E)** Identified motifs enriched within the ChIPmentation peaks from hCTCF-expressing yeast. Similarity to hCTCF motif is indicated in the third column.

**Supplementary Figure 3.**
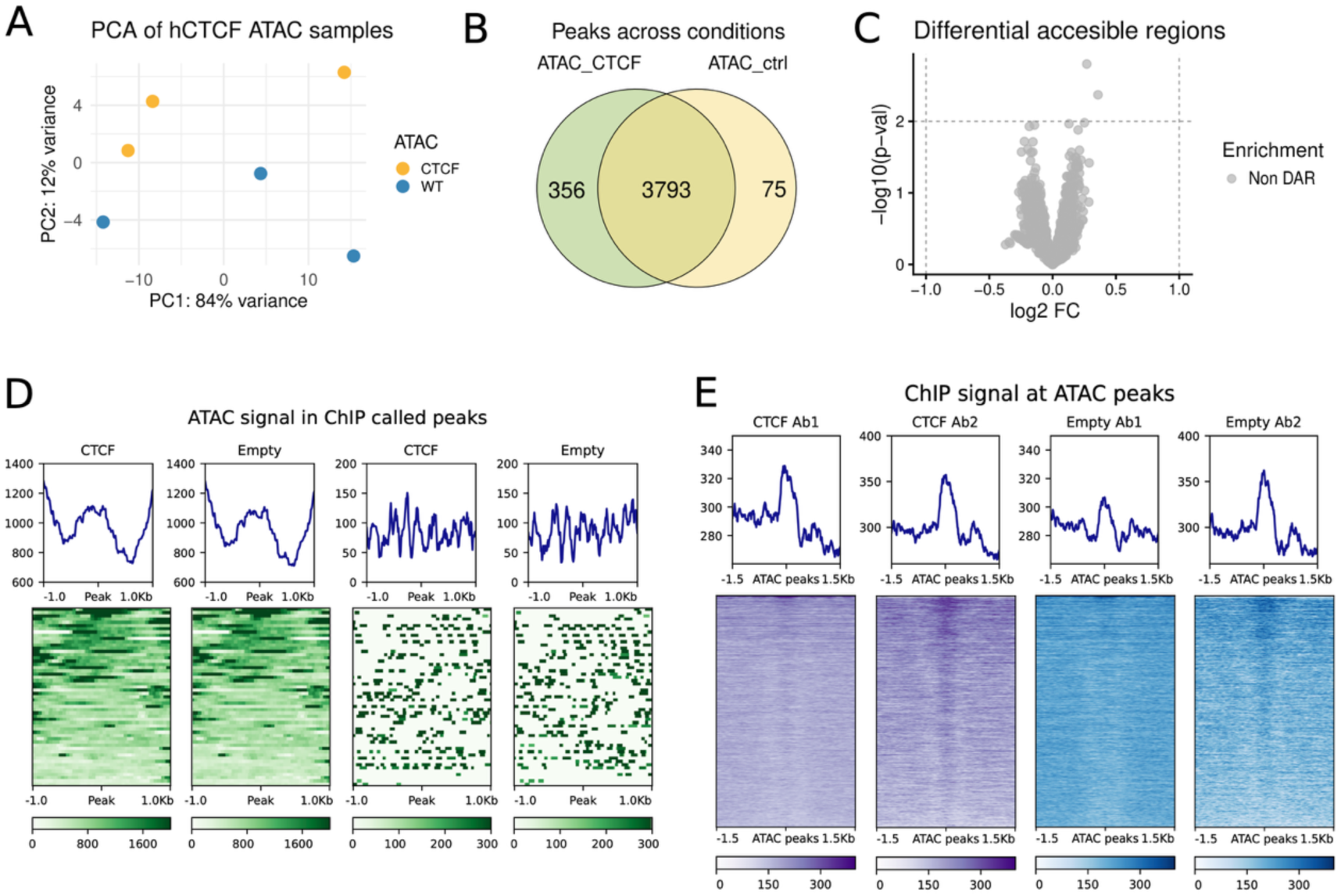
Chromatin accessibility after the expression of hCTCF. **(A)** Principal component analysis of ATAC-seq profiles from hCTCF expressing yeasts and control yeasts. Each point represents one biological replicate (n=3 per group). PC1 and PC2 explain 84% and 12% of the variance, respectively. **(B)** Venn diagram showing the overlap of the ATAC-seq peaks between conditions. **(C)** Volcano plots showing differential accessible regions (DARs) between hCTCF-expressing yeast and control dataset. **(D)** ATAC-seq signal profiles (left) and nucleosome occupancy (right) across a 2 kb region centered on hCTCF ChIPmentation peaks in hCTCF-expressing and control yeast. **(E)** ChIPmentation signal profiles across a 3 kb region centered on the ATAC-seq peaks identified in hCTCF-expressing yeast. From left to right, profiles correspond to hCTCF-expressing yeast (purple) using antibody 1 and antibody 2, followed by control yeast (blue) using antibody 1 and antibody 2.

