## Supplementary Figure 1 for "A naïve genome reveals the evolutionary dependence of CTCF functions"

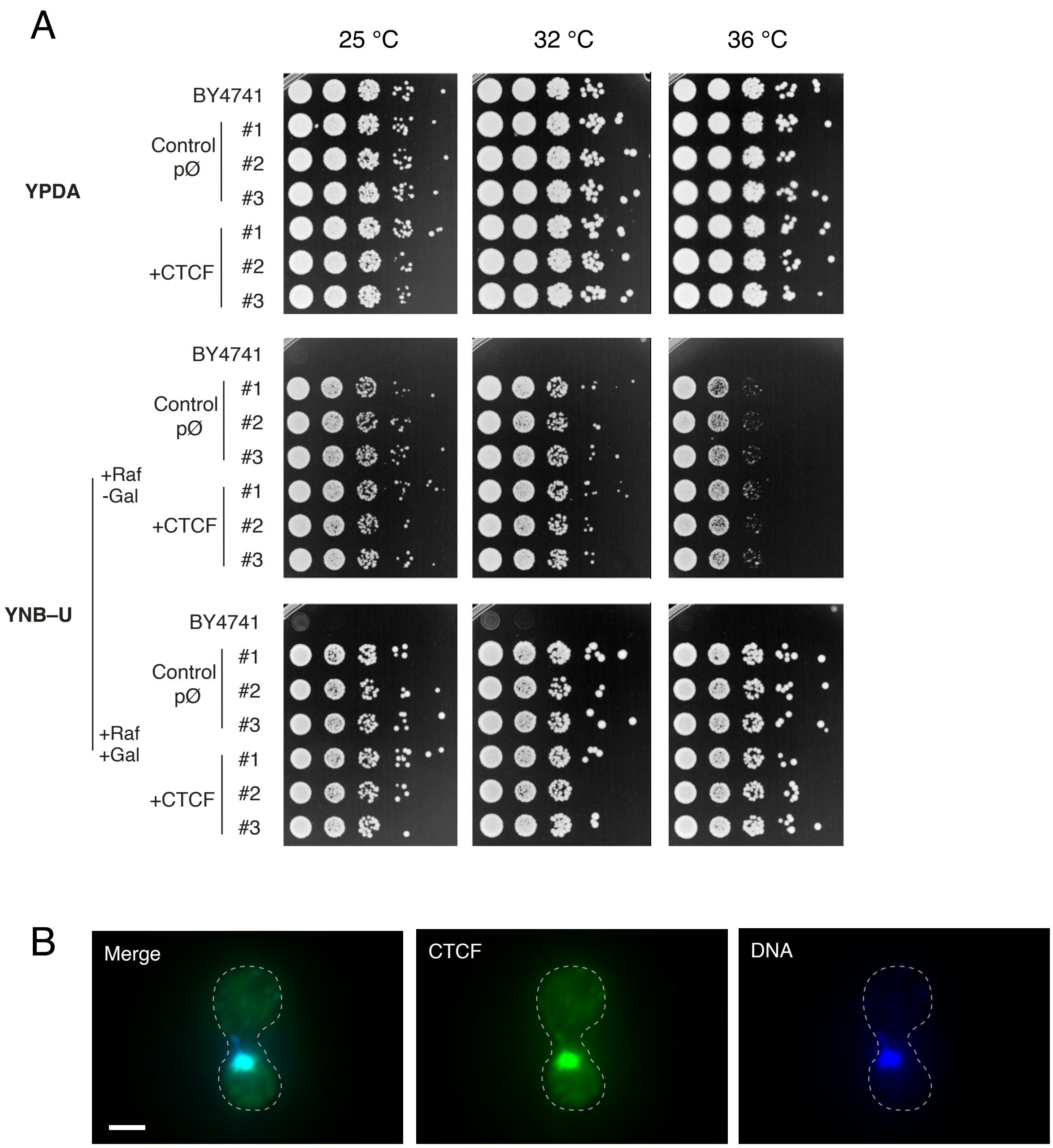
A
25 °C
32 °C
36 °C
YPDA
BY4741
Control
pØ
#1
#2
#3
+CTCF
#1
#2
#3
YNB–U
+Raf
-Gal
BY4741
Control
pØ
#1
#2
#3
+CTCF
#1
#2
#3
+Raf
+Gal
BY4741
Control
pØ
#1
#2
#3
+CTCF
#1
#2
#3
B
Merge
CTCF
DNA
