## Supplementary Figure 2 for "A naïve genome reveals the evolutionary dependence of CTCF functions"

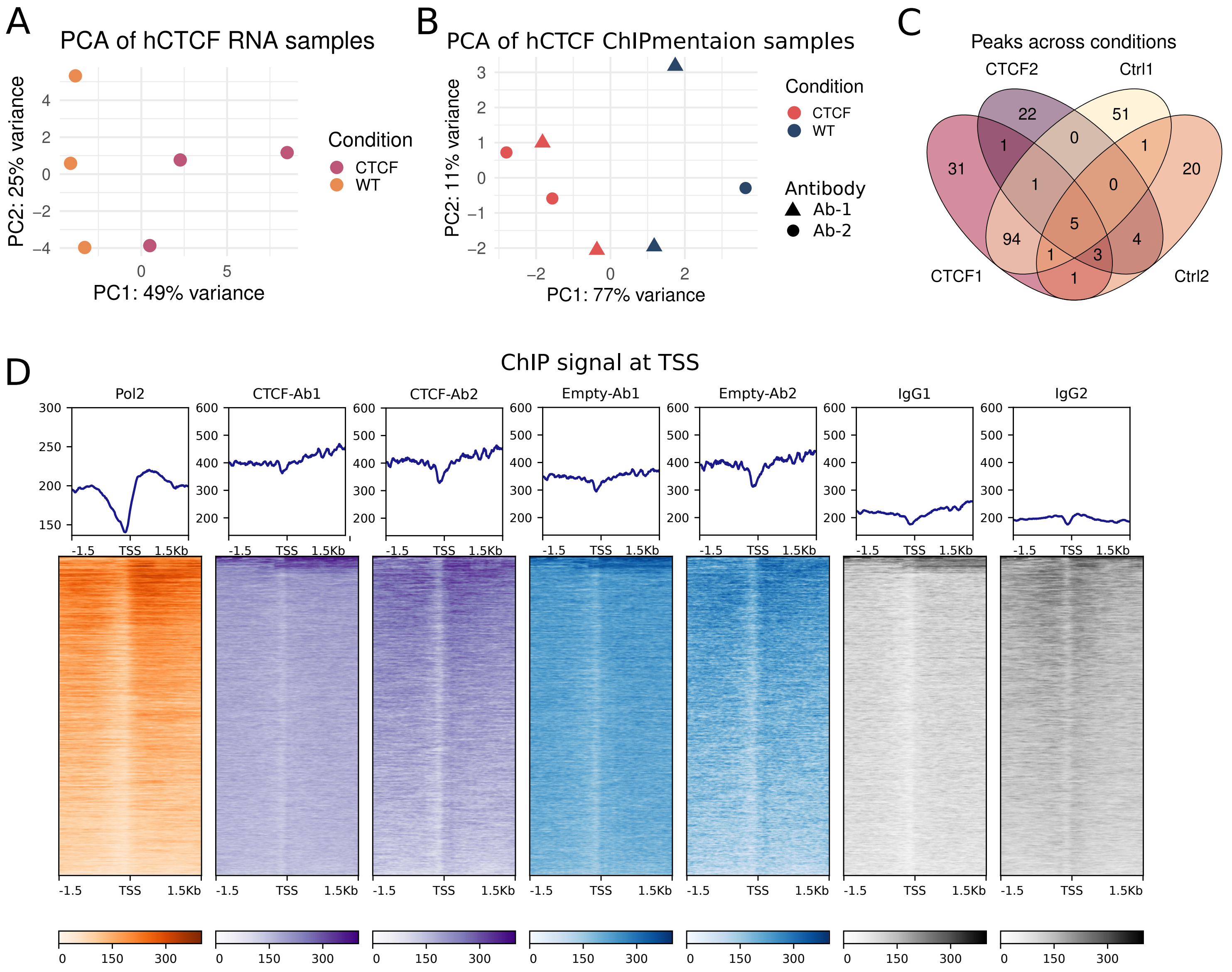


E

| Motif | Image | Similarity_to_CTCF | P-value | log10 P-value | ROC AUC |
|---|---|---|---|---|---|
| GimmeMotifs_12 | 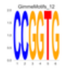 | 0.6659 | 6.98e-02 | 1.156 | 0.569 |
| GimmeMotifs_11 | 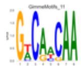 | 0.6492 | 2.89e-01 | 0.539 | 0.606 |
| GimmeMotifs_14 | 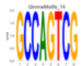 | 0.6492 | 4.92e-02 | 1.308 | 0.541 |
| GimmeMotifs_2 | 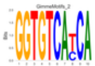 | 0.6463 | 1.26e-05 | 4.899 | 0.621 |
| GimmeMotifs_4 | 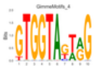 | 0.6463 | 3.60e-06 | 5.448 | 0.642 |
| GimmeMotifs_5 | 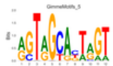 | 0.6419 | 4.11e-04 | 3.386 | 0.633 |
| GimmeMotifs_1 | 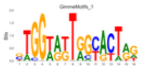 | 0.6148 | 2.00e-07 | 6.618 | 0.761 |
| GimmeMotifs_3 | 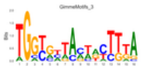 | 0.6148 | 0.00e+00 | 18.883 | 0.708 |
| GimmeMotifs_6 | 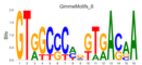 | 0.6148 | 0.00e+00 | 7.700 | 0.583 |
| GimmeMotifs_9 | 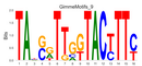 | 0.6148 | 0.00e+00 | 7.738 | 0.594 |
| GimmeMotifs_7 | 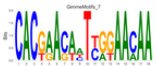 | 0.6021 | 4.00e-05 | 4.398 | 0.550 |
| GimmeMotifs_8 | 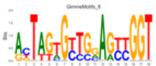 | 0.6021 | 3.64e-03 | 2.439 | 0.613 |
| GimmeMotifs_13 | 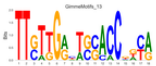 | 0.6021 | 0.00e+00 | 9.057 | 0.598 |
| GimmeMotifs_10 | 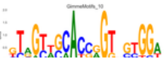 | 0.5953 | 0.00e+00 | 9.057 | 0.644 |
