## Supplementary Figure 3 for "A naïve genome reveals the evolutionary dependence of CTCF functions"

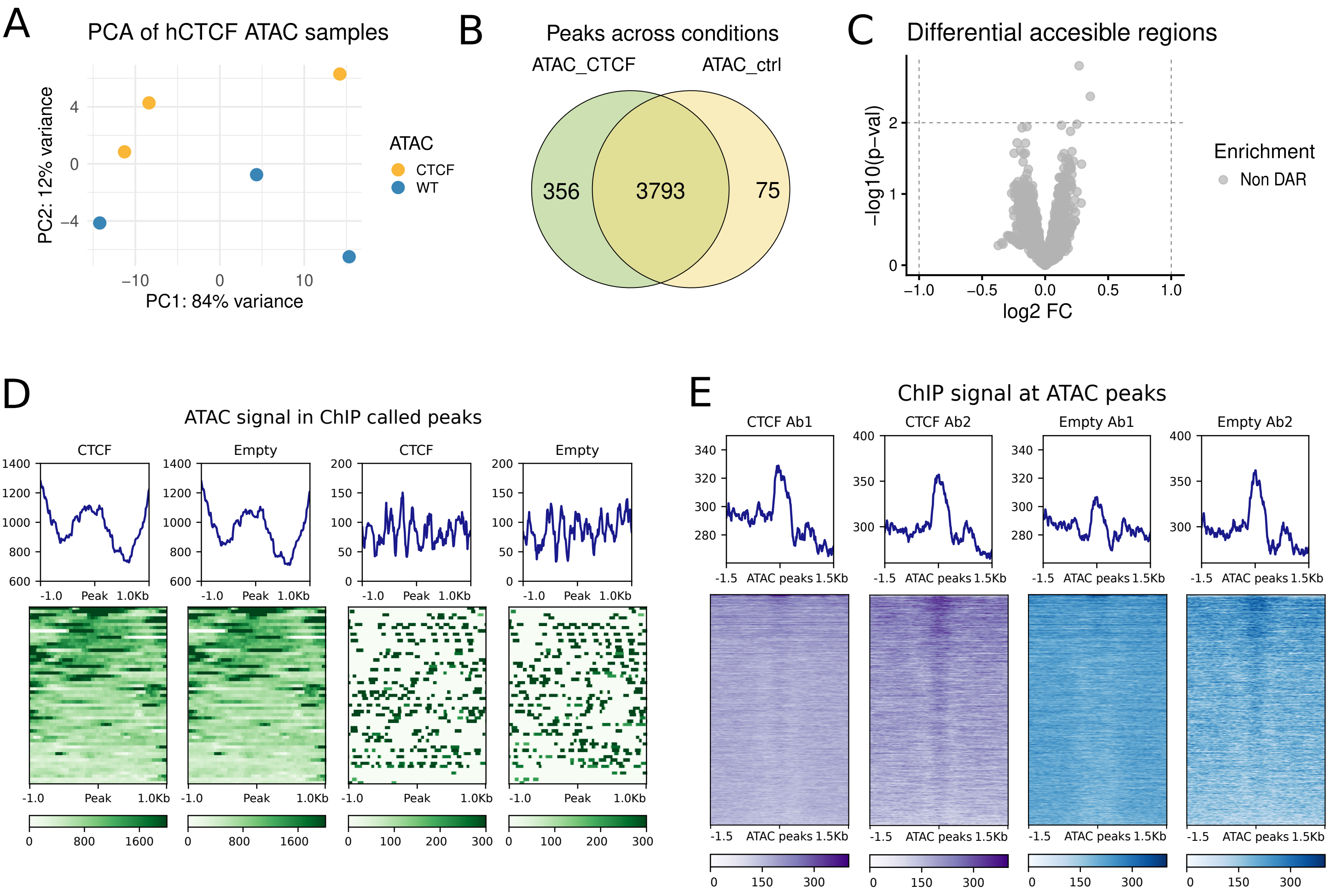
A
PCA of hCTCF ATAC samples
PC2: 12% variance
PC1: 84% variance
ATAC
CTCF
WT
B
Peaks across conditions
ATAC_CTCF
ATAC_ctrl
356
3793
75
C
Differential accesible regions
−log10(p−val)
log2 FC
Enrichment
Non DAR
D
ATAC signal in ChIP called peaks
CTCF
Empty
CTCF
Empty
E
ChIP signal at ATAC peaks
CTCF Ab1
CTCF Ab2
Empty Ab1
Empty Ab2
-1.5 ATAC peaks 1.5Kb
